# Variability in infants’ functional brain network connectivity is associated with differences in affect and behavior

**DOI:** 10.1101/2020.07.15.204271

**Authors:** Caroline M. Kelsey, Katrina Farris, Tobias Grossmann

## Abstract

Variability in functional brain network connectivity has been linked to individual differences in cognitive, affective, and behavioral traits in adults. However, little is known about the developmental origins of such brain-behavior correlations. The current study examined functional brain network connectivity and its link to behavioral temperament in newborn and 1-month-old infants (*M* [age] = 25 days; N = 75) using functional near-infrared spectroscopy (fNIRS). Specifically, we measured long-range connectivity between cortical regions approximating fronto-parietal, default mode, and homologous-interhemispheric networks. Our results show that connectivity in these functional brain networks varies across infants and maps onto individual differences in behavioral temperament. Specifically, connectivity in the fronto-parietal network was positively associated with regulation and orienting behaviors, whereas connectivity in the default mode network showed the opposite effect on these behaviors. Our analysis also revealed a significant positive association between the homologous-interhemispheric network and infants’ negative affect. The current results suggest that variability in long-range intra-hemispheric and cross-hemispheric functional connectivity between frontal, parietal and temporal cortex is associated with individual differences in affect and behavior. These findings shed new light on the brain origins of individual differences in early-emerging behavioral traits and thus represent a viable novel approach for investigating developmental trajectories in typical and atypical neurodevelopment.

---

Spontaneous brain activity is characterized by intrinsic dynamics of synchronized low-frequency fluctuations within structurally and functionally connected brain networks (Damoiseaux et al., 2006; Kaiser, Andrews-Hanna, Wager, & Pizzagalli, 2015). Much research has been focused on mapping the human connectome and delineating its anatomical and functional properties (Smith et al., 2013). Individual variability in functional connectivity profiles can accurately identify specific individuals likened to a fingerprint (Finn et al., 2015) and is linked to individual differences in cognitive, affective and behavioral traits in adults (Jauniaux, Khatibi, Rainville, & Jackson, 2019; Kaiser, Andrews-Hanna, Wager, et al., 2015; Picó-Pérez et al., 2019). More generally, the study of functional brain network connectivity has been argued to be one of the most promising and effective ways in bridging between brain and behavior (Friston, 2011).

From a developmental perspective, functional connectivity within brain networks can be detected from very early in human brain development. A host of studies employing resting-state functional magnetic resonance imaging (rs-fMRI) have mapped and identified functional networks in newborn infants (Fransson et al., 2009; Gao, Alcauter, Smith, Gilmore, & Lin, 2015; Schöpf, Kasprian, Brugger, & Prayer, 2012; Thomason et al., 2019; Zhang, Shen, & Lin, 2019). In fact, a body of work relying on progress in fetal rs-fMRI suggests that the basic organization and architecture of the functional connectome emerges during the late second trimester of pregnancy (Schöpf et al., 2012; Thomason et al., 2013; Thomason et al., 2019; van den Heuvel & Thomason, 2016). In addition, already within the first week of life early prenatal and postnatal experiences, such as premature birth, have begun to shape the development of these network (Rogers, Lean, Wheelock, & Smyser, 2018; Smyser, Wheelock, Limbrick, & Neil, 2019). The existing research with fetuses and infants points to a developmental progression whereby functional connectivity in primary short-range sensory-motor and homologous-interhemispheric networks are already in place at birth, whereas functional connectivity in higher-order cortical networks across longer ranges involving frontal, temporal and parietal cortex shows more protracted development during infancy (Gao et al., 2015; van den Heuvel & Thomason, 2016). For example, there is evidence to suggest that higher-order networks such as the default mode network (DMN) exist in a rudimentary form even in the fetus and newborn (Gao, Lin, Grewen, & Gilmore, 2017; van den Heuvel & Thomason, 2016), but functional network integration and synchronization continues to develop during infancy and beyond (Gilmore, Knickmeyer, & Gao, 2018). Other higher-order networks, such as the fronto-parietal network, show an even more protracted development as it is still considered immature by the end of the first year of postnatal life (see Gao et al., 2017, for a review). Moreover, it is these networks with the prolonged development, such as the fronto-parietal network, that go on to have the greatest inter-person variability and are considered to be unique identifiers of individuals (Finn et al., 2015). Taken together, much progress has been made in mapping the functional connectome in early human brain development, however, to date, little is known about whether and how functional brain network connectivity in these networks is linked to early affective, cognitive, and behavioral traits. This is a particularly important question considering that many mental health disorders are: (a) accompanied by alterations in functional connectivity (Hu et al., 2017; Kaiser, Andrews-Hanna, Wager, et al., 2015; Steward, Menchon, Jiménez-Murcia, Soriano-Mas, & Fernandez-Aranda, 2018) and (b) are argued to have deep developmental origins (Insel, 2010).

Depression as one main neuropsychiatric disorder affecting about 20.6% of the US population across their lifetime has been linked to specific alterations in functional brain network connectivity (Hasin et al., 2018). Specifically, a host of studies show that depression in adults is characterized by: (1) hypoconnectivity within the fronto-parietal network (FPN; composed of regions in the anterior cingulate cortex, dorsolateral prefrontal cortex, and parietal cortex) implicated in the cognitive control of attention and emotion (2) hyperconnectivity within the default mode network (DMN; composed of regions in the medial prefrontal cortex (mPFC), the precuneus, the posterior and anterior cingulate cortex, the inferior parietal cortex, and the lateral temporal cortex) involved in internally-oriented thought, mind-wandering, social cognition and (3) hypoconnectivity within the homologous-interhemispheric network (HIN; examining cross-hemispheric connections between frontal, temporal, and parietal lobes) involved in emotion regulation (Banich & Karol, 1992; Kaiser, Andrews-Hanna, Wager, et al., 2015; Patashov, Goldstein, & Balberg, 2019; Wang et al., 2013). Considering these alterations in functional connectivity seen in depression and the possibility that they might have their origins in early human brain development, it is important to examine variability in these brain networks and how it links to individual differences in affective and behavioral traits.

Therefore, the current study followed two major goals. First, we aimed to identify and map individual variability in the three functional brain networks (FPN, DMN, HIN) in young infants using functional near-infrared spectroscopy (fNIRS). FNIRS is a non-invasive, portable, and safe, optical neuroimaging technique for assessing functional connectivity in cortical brain networks during infancy (see the following papers for other examples of functional connectivity analysis using fNIRS with infants Bulgarelli et al., 2019; Bulgarelli et al., 2020; Homae et al., 2010). To capture functional connectivity patterns in young infants using fNIRS we pre-defined the following three long-range brain networks including available channels in specific frontal, temporal and parietal regions: 1) the FPN was created by measuring functional connectivity between the dorsolateral prefrontal cortex and inferior parietal cortex, 2) the DMN was created by measuring functional connectivity between the lateral temporal cortex and medial prefrontal cortex (note that our probe layout did not allow us to measure activity from superior parietal cortical regions including the precuneus, which is typically included in the DMN), and 3) the HIN was created by measuring functional connectivity between homologous cross-hemispheric connections in frontal, temporal, parietal cortex. In addition, based previous work measuring functional connectivity using fNIRS in adults (Sasai, Homae, Watanabe, & Taga, 2011), we created a so-called control network, computing functional connectivity between left frontal cortex and right temporal cortex and right frontal cortex and left temporal cortex. This served as a non-functional control network, because it is not known to have any functional associations and shows much lower levels of functional connectivity than established functional brain networks (Sasai et al., 2011).We hypothesized that functional connectivity within the three functional brain networks (FPN, DMN and HIN) will be significantly greater than in the control network, attesting to the existence of these long-range cortical networks in young infants. In this context, it is important to mention that, to our knowledge, there is no prior work demonstrating long-range functional connectivity in FPN and DMN in newborns and 1-month-old infants (Gao et al., 2017; van den Heuvel & Thomason, 2016), whereas functional connectivity in HIN has been shown to exist in the fetal brain (Thomason et al., 2013).

Critically, we also examined whether and how variability in functional brain network connectivity maps onto individual differences in infant temperament, which can be readily and reliably assed through parental report (Rothbart, 2007). Infant and child temperament is considered to reflect robust, biologically-based individual differences in affective and behavioral traits (Rothbart, 2007) linked to long-term developmental outcomes in prospective longitudinal studies, including adult personality, depression and anxiety (Asendorpf, Denissen, & van Aken, 2008; Caspi et al., 2003; Caspi, Moffitt, Newman, & Silva, 1996; Tang et al., 2020). In addition, brain networks assessed during the first few weeks of life has been linked to behavioral temperament 6 months later (Graham et al., 2016; Thomas et al., 2019). However, no work to the authors knowledge have examined the concurrent relation temperament in infants younger than 6 months. Specifically, we focused our investigation on three critical dimensions of infant temperament (regulation/orienting, negative emotionality, positive emotionality/surgency), which have been previously identified in a factor analysis (Gartstein & Rothbart, 2003). Based on prior work with adults linking functional connectivity in FPN to cognitive control of attention and behavior (Kaiser, Andrews-Hanna, Spielberg, et al., 2015; Kaiser, Andrews-Hanna, Wager, et al., 2015), we hypothesized that infants’ regulation/orienting behaviors will be associated with functional connectivity in the FPN with greater connectivity in this network being linked to enhanced regulation and orienting. In contrast, we predicted the opposite pattern of association in the DMN, lower connectivity being linked to enhanced regulation and orienting, as it has previously been linked to self-referential, stimulus-independent thought, and mind-wandering in adults (Kaiser, Andrews-Hanna, Spielberg, et al., 2015; Kaiser, Andrews-Hanna, Wager, et al., 2015). Moreover, we expected that infants’ reduced functional connectivity in the HIN will be linked to higher levels of negative emotionality, based on previous findings linking reduced cross-hemispheric connectivity to depression in adults (Patashov et al., 2019; Wang et al., 2013). Critically, in our analysis, we expected to see these predicted associations only for the specific functional networks but not for the (non-functional) control network. Finally, considering that there is no prior work informing how surgency/positive emotionality is linked to network connectivity, we did not have a specific hypothesis regarding this trait, but still included it in our analysis because surgency/positive emotionality has been identified as an important factor in previous work (Gartstein & Rothbart, 2003). Together, the current study presents a systematic examination of functional connectivity in long-range brain networks and its links to behavioral temperament in young infants.

## Materials and Methods

Seventy-five newborn and one-month-old infants (*M* [age] = 25 days; *Median* [age] = 24 days; ranging from 9 days to 56 days; 32 females; 43 males) were included in the final sample used for the present analyses. Participants were recruited from a local hospital. The diverse sample of infants were representative of the surrounding Mid-Atlantic college town area such that the majority of infants were Caucasian (*n* = 49 Caucasian; *n* = 14 Black; *n* = 3 South Asian; *n* = 3 Pacific Islander; *n* = 2 Asian; *n* = 4 Other), from highly-educated parents (*n* = 31 obtained a Graduate Degree; *n* = 19 Bachelor’s Degree; *n* = 12 some College/Associates Degree; *n* = 11 High School Diploma/GED; *n* = 2 some High School), and low to medium-income families (*n* = 21 $15-45,000; *n* = 18 $75-110,000; *n* = 11 $45-75,000; *n* = 11 $110-175,000; *n* = 8 $175,000+; *n* = 5 less than $15,000; *n* = 1 did not respond). All participants were born at term, with normal birth weight (>2,500g), and did not have any hearing or visual impairments. Thirty-three additional infants were tested but were excluded from the present analyses for the following reasons: *n* = 25 were excluded because they failed to reach our pre-determined inclusion criterion of having at least 100 seconds of continuous data with non-disruptive behaviors (see below); *n* = 4 were excluded because of inaccurate placement of the cap; *n* = 4 were excluded because more than 33% of the measured fNIRS channels had poor light intensity readings, more specifically, a signal-to-noise ratio of less than 1.5 (Bulgarelli et al., 2019; Xu et al., 2015). Note that the current attrition rate (30%) is lower than in previous infant fNIRS studies (Cristia et al., 2013). Moreover, temperament profiles (negative emotionality, regulation/orienting, and surgency/positive emotionality) were compared using independent samples t-tests between infants that were included and excluded from the present analyses and no significant differences were found between the two groups (all *p-values* > .29). All parents gave informed consent for their infants to participate in accordance with the Declaration of Helsinki and families received a payment for their participation. All procedures were approved by and carried out in accordance with The University of Virginia Institutional Review Board for Health Sciences (Protocol number 20381).

### Infant Temperament

Infant temperament was assed using parental reports of the 91-item Infant Behavior Questionnaire Revised Short Form (IBQ-R; Gartstein & Rothbart, 2003). Parents filled out the questionnaire online using Qualtrics survey platform prior to their appointment. This measure has been widely used and shown to be reliable and valid at the newborn time point (see the following papers for examples of prior work using this measure with newborns Rigato, Stets, Bonneville-Roussy, & Holmboe, 2018; Stifter & Fox, 1990; Worobey & Blajda, 1989). The questionnaire asks parents to report their infant’s behavior during the previous two weeks and rate the occurrence/frequency of the behavior on a 1 (Never) to 7 (Always) scale. Based on prior work using factor analysis (Gartstein & Rothbart, 2003), three general temperament dimensions were computed summarizing information from various sub-scales: (1) negative emotionality (contributing sub-scales: fear, distress to limitations, falling reactivity, sadness), (2) regulation/orienting (contributing sub-scales: low intensity pleasure, cuddliness, duration of orienting, soothability), and (3) surgency/positive emotionality (contributing sub-scales: activity level, smiling and laughing, high intensity pleasure, perceptual sensitivity, approach, vocal reactivity) (Gartstein & Rothbart, 2003). If parents reported the behavior was not applicable at the current time then this item was given a value of 0. Chronbach’s alpha coefficients were calculated to determine reliability of the temperament measures and all values were in acceptable ranges for each of the three dimensions: surgency/positive emotionality α = .78, regulation/orienting α = .78, and negative emotionality α = .91. Finally, correlation analyses between Edinburgh Postnatal Depression Scale scores (assessed at the same time as behavioral temperament; Cox, Holden, & Sagovsky, 1987) and behavioral temperament scores were conducted in order to statistically account for any variance in maternal-reported behavioral temperament that may be related to maternal mental health. Here, we did not find any significant associations (all *p-values* > .24). Therefore, maternal depression was not used as a covariate in later analyses.

### Procedure

The resting state fNIRS task took place in a quiet, dimly-lit testing area. Infants were seated on their parents’ lap and placed approximately 60 cm from the screen (23-inch monitor). The infants were fitted with a fNIRS fabric cap (EasyCap, Germany) which was secured in place using infant overalls and outside netting. The experimental paradigm was presented using the Presentation software package (Neurobehavioral Systems, USA). A non-social stimulus was created by selecting non-social clips from a popular infant video (Baby Einstein - Kids2 Inc.) that featured videos of toys, stuffed animals, and still images of everyday objects, which was accompanied by classical music (Lordier et al., 2019). Similar screen-saver-like videos have been used in prior work examining functional connectivity using fNIRS (see Bulgarelli et al., 2020). This video was played for a total of seven minutes while fNIRS data were being recorded. The clips were segmented into 30 second intervals and the order of presentation was randomized for each infant. Parents were asked to remain quiet throughout the fNIRS recording session. Sessions were video-recorded using a camera mounted above the screen. This allowed for later offline coding of infants’ alertness and cap placement.

### Data acquisition

Infants’ fNIRS data were recorded using a NIRx Nirscout system and NirStar acquisition software. The fNIRS method quantifies concentration changes of oxygenated hemoglobin (oxyHb) and deoxygenated hemoglobin (deoxyHb) in the cerebral cortex through shining specific frequencies of light that are selectively absorbed by these chromophores (for more information regarding this technique see Lloyd-Fox, Széplaki-Köllőd, Yin, & Csibra, 2015). The fNIRS system used contains 16 source-detector pairs (approximately 2.0 cm apart) resulting in a total of 49 channels positioned over frontal and temporal-parietal regions (see Altvater-Mackensen & Grossmann, 2016; Grossmann, Missana, & Krol, 2018; Kelsey, Krol, Kret, & Grossmann, 2019; Krol, Puglia, Morris, Connelly, & Grossmann, 2019 for infant work using the identical channel positioning/layout). The system emits two wavelengths of light in the Near-Infrared spectrum, 760 nm and 850 nm, and captures both deoxyHb and oxyHb. The diodes have a power of 25 mW/wavelength and data were recorded at a preset default sampling rate of 3.91 Hz.

### Behavioral Coding

Infants’ behavior during the fNIRS recording session was coded by a trained research assistant using video recordings of the experimental session. Specifically, coders identified timepoints where the infants were crying, excessively moving, looking at the parents, or parents were talking in order to remove these periods from the analysis. To assess the reliability of the attentional coding done by the primary coder, an additional trained coder also coded infant behavior from selected subsample of infants (25.3%; *n*□=□19). This analysis showed that inter-rater reliability amount of data included was excellent (Cronbach’s α□=□0.94). In line with previous studies, infants were only included in the present analysis if they had at least 100 seconds of disruption-free (see aforementioned behaviors) data (Bulgarelli et al., 2019). Moreover, as it takes a minimum of 8 seconds for the Hemodynamic response function to return to baseline after a stimulus-evoked event, the onset of useable data was delayed for 8 seconds (Bulgarelli et al., 2019; Bulgarelli et al., 2020). However, unlike Bulgarelli et al. (2019; 2020), the time series of fNIRS obtained in the current study was continuous. On average, infants contributed 317.59 seconds of data (*SD* = 115.46 seconds; range = 100-420 seconds). Furthermore, the amount of data included in the current analysis is comparable to other functional connectivity work using fNIRS with older infants (Bulgarelli et al., 2020). In addition, we coded infants’ state of alertness on a 1 (Deep Sleep) to 6 (Crying) scale. On average infants were rated as being in a Active Light Sleep to Drowsy State (*M* = 2.68, *SD* = 1.27). Finally, we assessed how infants’ behavior throughout the session, specifically the amount of useable data related to functional network connectivity in each of the networks (see supplemental materials).

### Data Analysis

The fNIRS data were analyzed using the functional connectivity program, FC-NIRS (Xu et al., 2015). First, channels were assessed for light intensity quality and channels were removed if the signal-to-noise ratio was less than 1.5 (Xu et al., 2015). In order to be included in the present analyses, infants needed to have at least 70% of their channels passing this threshold (Bulgarelli et al., 2019). Next, data were band-pass filtered (using a .08 Hz low-pass filter, to remove fast fluctuations related to heart rate, and a high-pass filter of .01 Hz, to remove changes that were too slow and related to drift; Bulgarelli et al., 2019; Lu et al., 2009). This range of .01 to .08 Hz was chosen on the basis of prior work (Bulgarelli et al., 2019; Sasai et al., 2011). This range was also selected because it falls well below the reported range for cardiac fluctuations (greater than 1 Hz), providing us with greater confidence that the measured changes reflect hemodynamic events tied to cortical activity rather than (systemic) cardiovascular system activity (e.g., heart rate Elwell, Springett, Hillman, & Delpy, 1999; Obrig et al., 2000). Finally, concentration changes were calculated using the modified Beer-Lambert law [partial path length factor: 6.0] (Villringer & Chance, 1997).

For each infant, we obtained a 49 by 49 correlation matrix corresponding to all of the relations between all of the channels measured. Considering that negative values are difficult to interpret in terms of their neurobiological basis, and based on prior work, we first checked data to see if there were any negative values and found out there were none (Fox, Zhang, Snyder, & Raichle, 2009; Murphy, Birn, Handwerker, Jones, & Bandettini, 2009). In order to standardize the values, Fisher Z-transformations were performed on all correlation matrices. Networks of interest were created by selecting channels that corresponded to specific regions of interest. Brain networks were composed based on the anatomical information available in Kabdebon et al. (2014), a meta-analysis of resting state fMRI (Kaiser, Andrews-Hanna, Wager, et al., 2015), a large resting state fMRI functional connectivity analysis of newborn infants (Eyre et al., 2020), and prior work infant and adult work using rs-fNIRS (Bulgarelli et al., 2020; Imai et al., 2014; Patashov et al., 2019; Sasai et al., 2011). Based on this information four networks were created: (1) The FPN was created by averaging all correlations between three channels in the dorsolateral prefrontal cortex (corresponding with the F3, F4, F5, F6 electrodes) and two channels in the parietal area (corresponding with CP3 and CP4 electrodes); (2) The DMN was created by averaging all correlations between three channels in the medial prefrontal cortex (corresponding with the Fpz electrode) and four channels in the lateral temporal cortex (corresponding with FT7, T7, FT8, T8 electrodes); (3) The HIN was created by averaging all correlations between the 21 channels in the left hemisphere (including frontal, temporal and parietal cortical regions) with their corresponding (homologous) channels in the right hemisphere; and, (4) a (non-functional) control network was created by averaging all correlations between three channels in the left frontal area (corresponding with the F7 electrode) with three channels in the right temporal area (corresponding with the T8 electrode) and three channels in the right frontal area (corresponding with F8 electrode) with three channels in the left temporal area (corresponding with the T7 electrode; see Figure 1 for schematic of network configurations). Cortical projections onto a standard MNI newborn (0-2 months old) atlas (Fonov et al., 2011) were created using NIRSite (Nirx) by using 10-20 system references from the cap layout.

**Figure 1.**
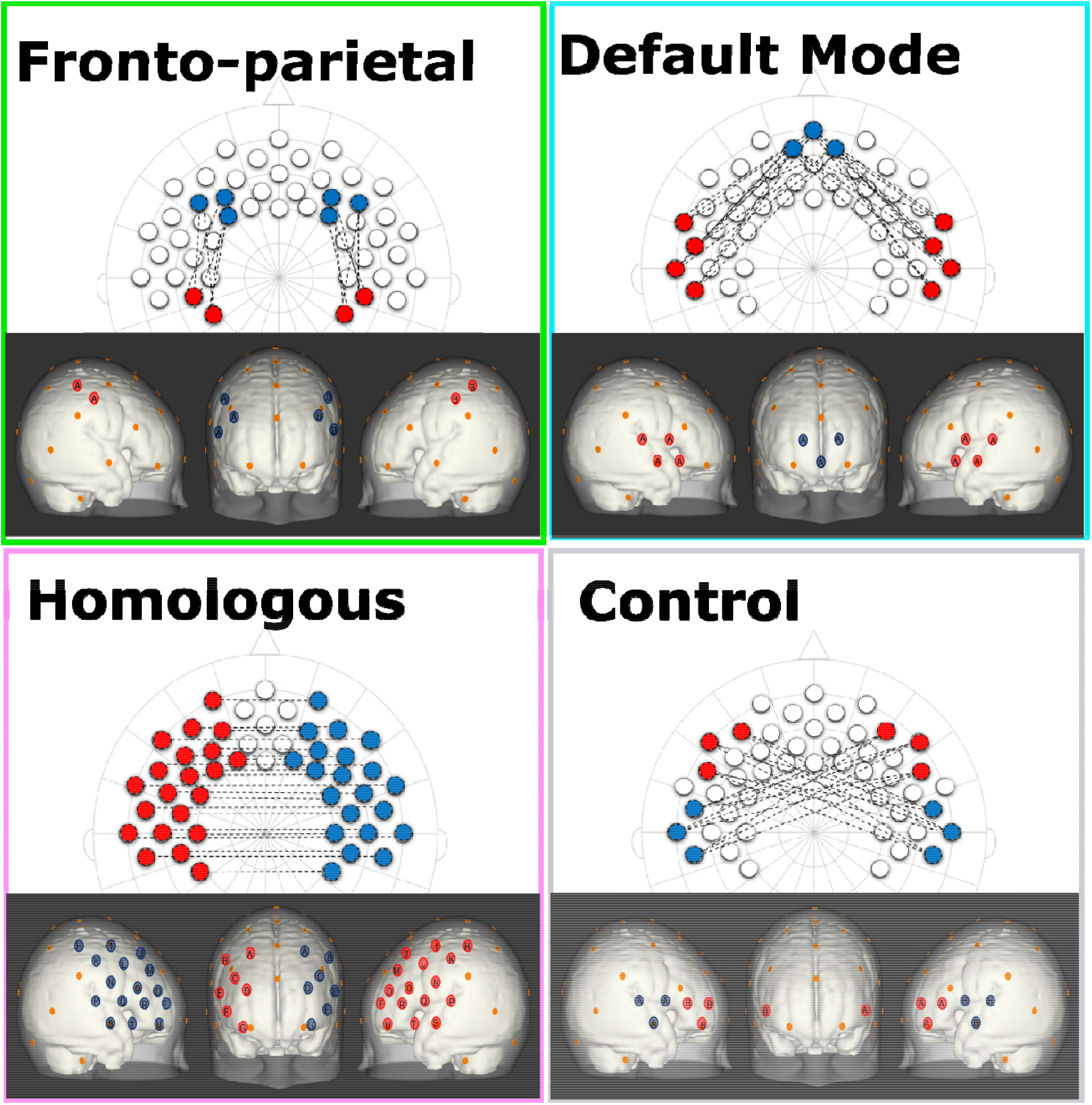
Shows the configurations for each of the network patterns in both a 2-dimensional 10-20 system layout and estimated projections onto cortical space of a 0-2 month-old Atlas (Fonov et al., 2011). Note, each network consists of the average of all of the connections between red and blue channels of the same letter. In addition, the orange dots represent relevant 10-20 landmarks.

All analyses were conducted for both oxyHb and deoxyHb (for deoxyHb results please see supplemental materials). Moreover, statistical outliers – values that were more than 3 SD above the mean – were removed for the subsequent analyses (FPN *n* = 2, negative emotionality *n* = 1).

## Results

A series of Spearman’s rho correlations were used to identify significant associations between variables of interest and potential socio-demographic factors (for a schematic representation for all associations see supplementary Figure 1). Any demographic variables found to be significantly associated with a study variable of interest were then included in the subsequent models assessing differences in said study variable as a covariate. Negative Emotionality was significantly associated with both infant age (Spearman’s rho correlation r_s_ = .47, *p* < .001) and family income (Spearman’s rho correlation r_s_= .29, *p* = .011). Regulation/orienting was significantly associated with Education (Spearman’s rho correlation r_s_= −30, *p* = .009). However, there were no significant associations found between any of the functional connectivity measures and any of the covariates.

### Functional connectivity across networks

As a first step, a series of one-sample t-tests were conducted to assess whether Fisher-transformed correlation between individual channels within the pre-defined networks of interest differed from zero. As shown in Figure 2, this analysis identified significant functional connectivity between individual channels within the pre-defined networks of interest (see supplementary Table 1). Next, we conducted a series of one-sample t-tests to assess connectivity at the network level (combining across all channels of interest). Here, all networks (FPN, DMN, HIN, control) were found to be greater than zero (all p’s < .001; see Figure 3). To analyze differences in overall connectivity levels across networks an omnibus repeated measures ANOVA with network type (FPN, DMN, HIN, control) as a within-subjects factor was conducted. This analysis revealed a significant within-subjects effect across network types, *F*(3, 216) = 18.78, *p* < .001, η^2^ = .207. Post-hoc analyses with Bonferroni adjustments for multiple comparisons were conducted to assess which networks significantly differed from one another. Importantly, all functional networks of interest had significantly higher connectivity than the (non-functional) control network (*M* = .05; *SD* = .12; range: −.20–.44), all p’s < .001. In addition, we found that there was significantly greater connectivity in the FPN (*M* = .21; *SD* = .20; range: −.16–.72) compared to both the HIN (*M* = .13; *SD* = .12; range: −.12–.48), *p* = .003, and the DMN (*M* = .13; *SD* = .16; range: −.28–.73), *p* = .01. However, there was no significant difference found between the level of connectivity for the HIN from the DMN, *p* = 1.00 (see Figure 3).

**Figure 2.**
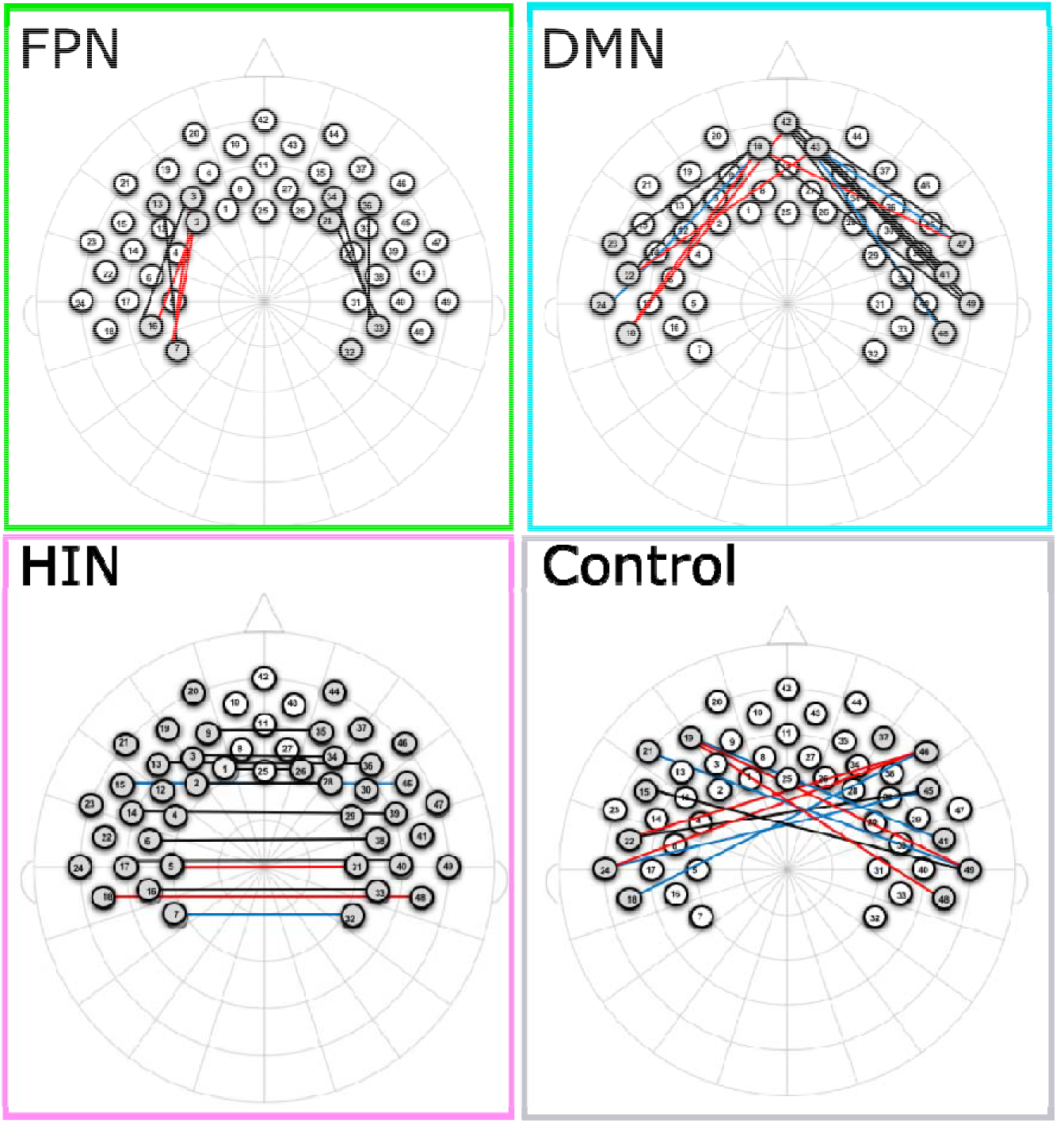
Shows the channels that are significantly different than zero for each of the networks. Please note, channels in red, blue, and black represent significant changes for oxyHb, deoxyHb, and both oxy and deoxyHb respectively.

**Figure 3.**
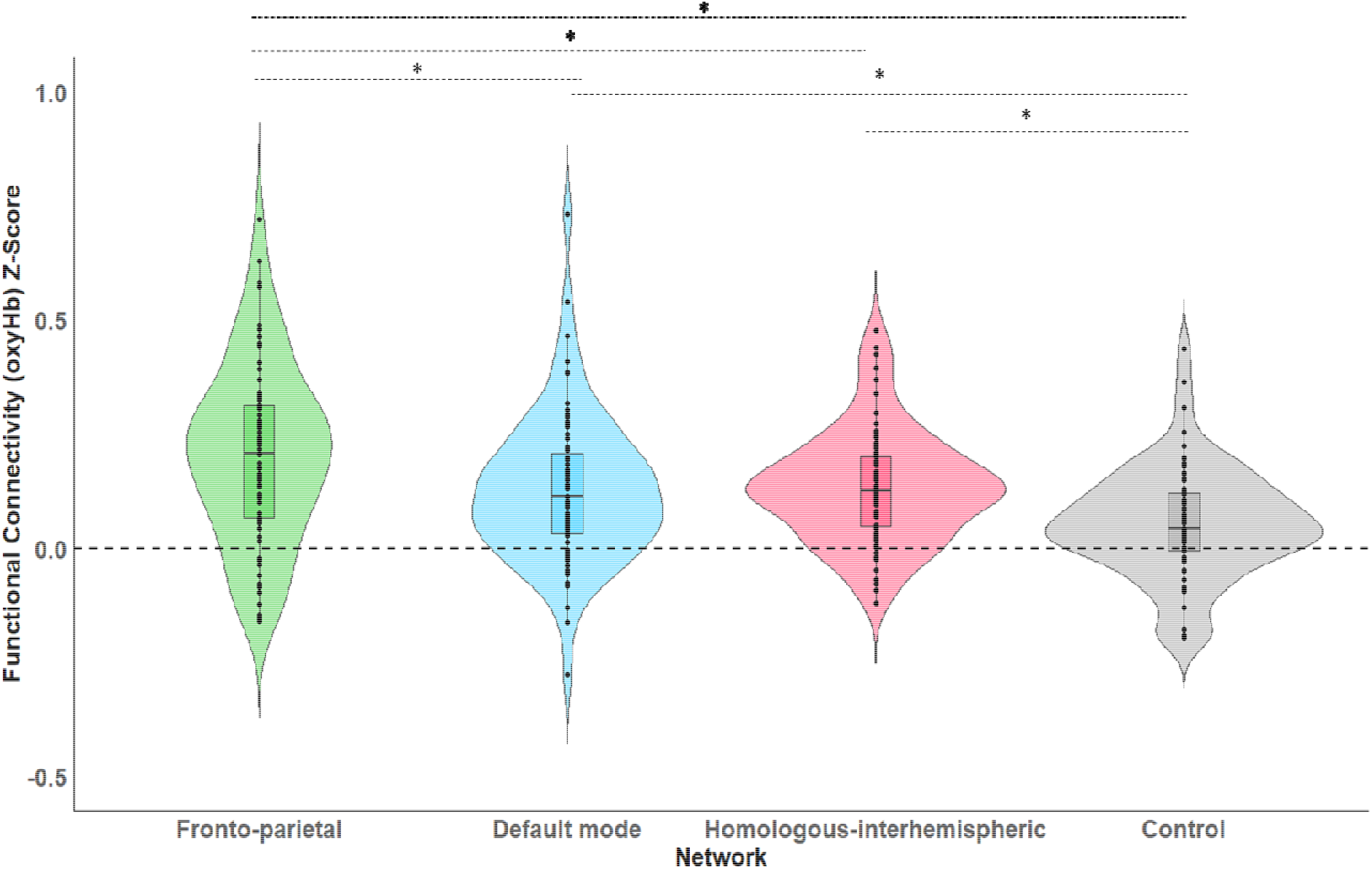
Shows the average levels of functional connectivity (oxyHb) and range of variability for each network. The boxplot horizontal lines from bottom to top reflect values for the lower quartile, median, and upper quartile respectively. Note, * *p* < .05.

### Functional connectivity and behavioral temperament

In order to assess how functional connectivity patterns differentially predicted temperament characteristics, three separate regressions with all four network types (FPN, DMN, HIN, control) predicting each of the three domains of temperament (negative emotionality, regulation/orienting, surgency/positive emotionality) were conducted.

#### Regulation/Orienting

A linear regression was conducted with the four network types (FPN, DMN, HIN, control) predicting regulation/orienting using the entry method. The regression model significantly predicted regulation/orienting, *F*(5, 72) = 4.84, *p* = .001, *R^2^* = .27. More specifically, connectivity in the DMN was negatively associated with regulation/orienting (β = −1.02, *SE* = .42, *p* = .018); whereas, connectivity in the FPN was positively associated with regulation/orienting (β = .71, *SE* = .35, *p* = .049; see Figure 4). Neither the HIN nor the Control network were found to be related to regulation/orienting, all *p-values* > .24.

**Figure 4.**
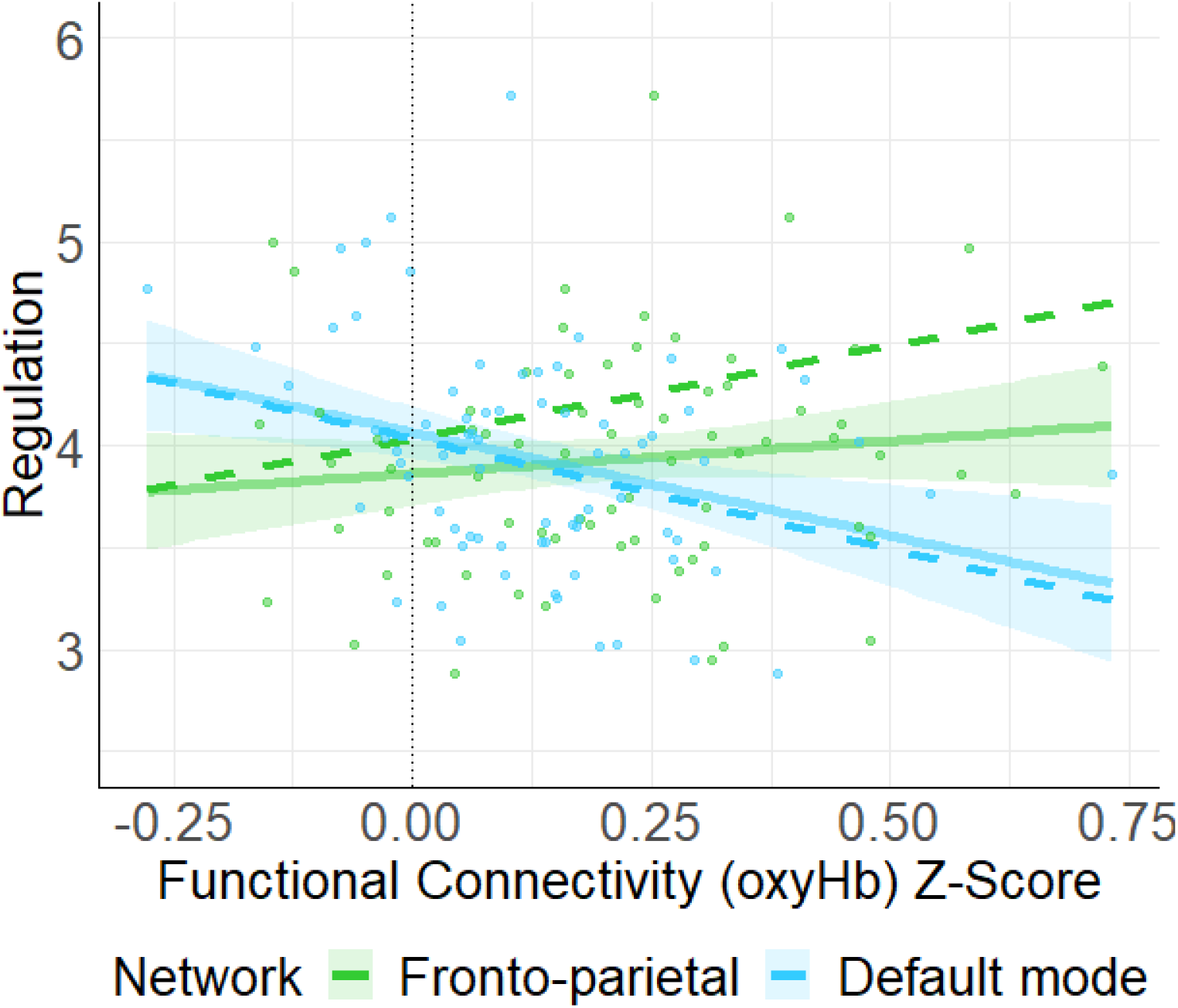
Shows the unadjusted (solid line) and adjusted relation (covariates not shown: HIN, Control) between functional connectivity (oxyHb) Z-score and regulation/orienting. Here, we found that connectivity in the FPN was positively associated with regulation/orienting (*p* = .049) whereas, connectivity in the DMN was negatively associated with regulation/orienting (*p* = .018). Note, shaded regions represent 90% confidence intervals for the raw (unadjusted) data.

#### Negative Emotionality

A multiple linear regression was conducted with the socio-demographic covariates (age, income) and four network types (FPN, DMN, HIN, control) predicting negative emotionality using the entry method. The model significantly predicted negative emotionality *F*(6, 64) = 5.50, *p* < .001, R^2^ = .34. More specifically, we found a significant positive relation between HIN connectivity and negative emotionality, (β = 1.60, *SE* = .75, *p* = .038; See Figure 5). However, none of the other networks (functional nor control) were found to be related to Negative Emotionality, all *p-values* > .57.

**Figure 5.**
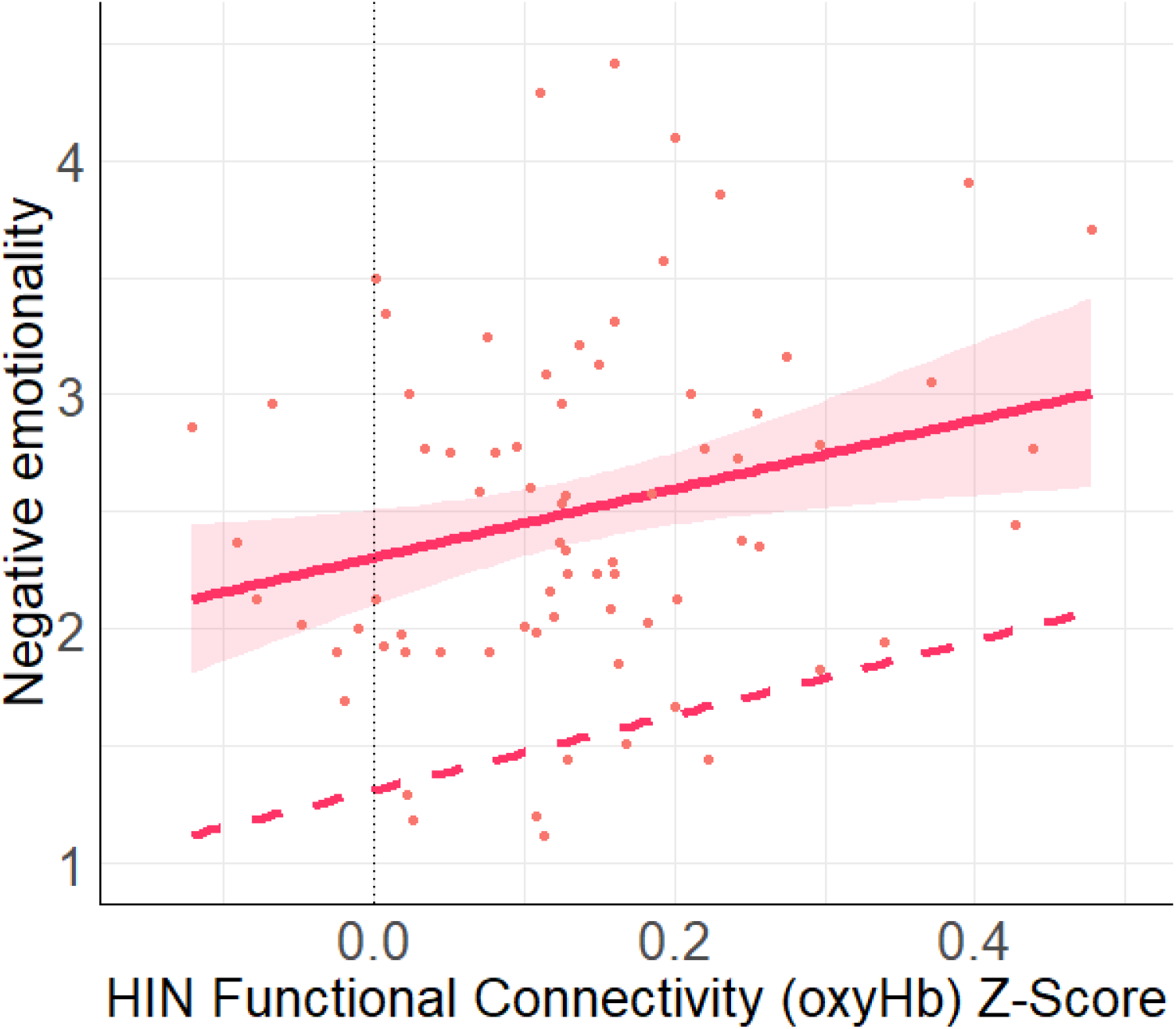
Shows the unadjusted (solid line) and adjusted relation (covariates: FPN, DMN, Control, Income, age) between HIN functional connectivity (oxyHb) Z-score and negative emotionality. Here, we found a significant positive relation between the HIN and negative emotionality (*p* = .038). Note, shaded regions represent 90% confidence interval for the raw (unadjusted) data.

#### Surgency/Positive Emotionality

A linear regression was conducted with the four network types (FPN, DMN, HIN, control) predicting surgency/positive emotionality using the entry method. Here, the regression model did not significantly predict surgency/positive emotionality, *p* = .39. Moreover, none of the network types were significantly associated with surgency/positive emotionality (all *p*’s > .14).

## Discussion

The current study examined functional connectivity in brain networks using fNIRS and behavioral temperament using parental report in young infants. Our results show that functional connectivity in long-range cortical brain networks (FPN, DMN and HIN) can be identified in very young infants and that functional connectivity in these networks varied considerably among infants. This supports the suitability of fNIRS in assessing functional connectivity and its variability in newborn infants. Importantly, our results also show that such variability in functional brain network connectivity systematically maps onto individual differences in infant behavioral temperament. Overall, the current findings provide novel insights into the brain origins of individual differences in affect and behavior, pointing to the early perinatal foundation of human temperament.

In line with our hypothesis, functional connectivity within the three brain networks (FPN, DMN, and HIN) was significantly greater than in the control network and significantly greater than a zero-value, indicating the existence of these long-range cortical brain networks in young infants. This provides further evidence that functional brain networks exist from early in ontogeny and are detectable in young infants (Gao et al., 2017; Gao et al., 2009; Zhang et al., 2019). To our knowledge, this is the first study to demonstrate long-range functional connectivity in FPN and DMN in young infants, suggesting a remarkably early emergence of long-range connectivity in higher-order brain networks linked to cognitive control and self-referential processes, respectively. The current findings are noteworthy also in regard to the fact that both networks involve regions in prefrontal cortex, providing new evidence from newborns and 1-month-old infants supporting the view that prefrontal cortex plays a critical role in human brain function from very early in development (Grossmann, 2013a, 2013b; Grossmann, 2015; Powell, Kosakowski, & Saxe, 2018; van den Heuvel & Thomason, 2016).

In addition to the general difference in connectivity between the functional and the (non-functional) control network, we also found that activity in the FPN was significantly greater than in the DMN and HIN (whereas there was no difference in connectivity levels found between the DMN and HIN). One possible interpretation of this finding is that functional connectivity in the FPN might have been enhanced when compared to the other functional networks, because like other resting-state studies with infants, the participants were presented with a video accompanied by music during the fNIRS measurement (Bulgarelli et al., 2019). In other words, the FPN might have been more engaged because infants were attending to external audio-visual stimuli (note that all infants were exposed to the same video [audio-visual] stimulus). Here it is important to mention that prior work with adults using fMRI shows that functional connectivity in higher-order cortical resting-state networks can be reliably acquired during the presentation of videos and corresponds to functional connectivity acquired in the absence of any stimulus (Finn et al., 2015; Vanderwal, Kelly, Eilbott, Mayes, & Castellanos, 2015). Nonetheless, based on recent work showing that preterm infants display enhanced functional connectivity in higher-order cognitive networks in response to music (Lordier et al., 2019), we speculate that enhanced functional connectivity in FPN might at least be partly explained by having newborn infants listen to music in the current study. Clearly, future research with infants, which systematically compares between stimulation protocols, is needed to examine whether and how functional connectivity is influenced by the measurement context and the stimulation protocol used. Overall, our functional connectivity analysis supports the notion that intrinsic functional connectivity in cortical brain networks and its variability can be effectively mapped in newborn infants using fNIRS.

Having established functional connectivity in these brain networks as variable and distinct from a (non-functional) control network then allowed for the examination of specific associations between brain network connectivity and infant behavioral temperament. Our results confirmed our hypothesis and showed that infants’ regulation/orienting behaviors were associated with functional connectivity in the FPN with greater connectivity in this network being associated with enhanced regulation and orienting. This result is in line with prior work linking functional connectivity in FPN to cognitive control of attention and behavior in adults (Kaiser, Andrews-Hanna, Spielberg, et al., 2015; Kaiser, Andrews-Hanna, Wager, et al., 2015). The current results further showed the opposite pattern of association for functional connectivity in DMN, with greater connectivity associated with reduced regulation and orienting, which is in agreement with our hypothesis based on the DMN previously being linked to self-referential, stimulus-independent thought and mind-wandering in adults (Kaiser, Andrews-Hanna, Spielberg, et al., 2015; Kaiser, Andrews-Hanna, Wager, et al., 2015). To obtain such opposing effects of functional connectivity in FPN and DMN is reminiscent of seminal findings supporting the existence of anti-correlated brain networks in adults (Fox et al., 2005) and may suggest that similar organizational principles are at play in newborn infants. However, it should be emphasized that, functional connectivity in FPN and DMN in the current study was not anticorrelated as such, but rather had opposing effects on infants’ behavioral and attentional regulation.

Assuming that depression represents variability at more extreme ends of a spectrum in FPN and DMN function, our results concerning behavioral and attentional regulation and their functional connectivity correlates in infants, are principally in line with prior research showing hyperconnectivity in the DMN and hypoconnectivity in the FPN in adults with depression (Kaiser, Andrews-Hanna, Wager, et al., 2015). Moreover, our data show that infants’ functional connectivity in the HIN was associated with negative emotionality. Contrary to prior work with adults indicating that hypoconnectivity is associated with depression (Patashov et al., 2019; Wang et al., 2013), the current infant data show that greater connectivity between homologous brain regions in both hemispheres was associated with greater negative affect. It is unclear why the direction of the association (positive versus negative) would differ as a function of age (newborn infants in the current study and adults in previous work), but it is worth noting that the experience and display of negative affect only gradually emerges during infancy and may thus not be fully present in newborn infants (Stifter & Fox, 1990). The current approach of measuring functional connectivity in brain networks during early development previously shown to be associated with depression therefore affords the possibility in designing protocols to assess risk for developing depression. Future research employing prospective longitudinal designs is needed to systematically investigate these brain-behavior correlations with implications for our understanding and treatment of depression.

Taken together, the current findings demonstrate specific associations between functional brain network connectivity and behavioral temperament in newborn infants. This suggests a remarkably early emergence of functional networks with behavioral relevance and highlights the importance of evaluating individual differences reflected in intrinsic brain connectivity. Although there are many advantages in the current approach of using fNIRS to examine functional brain connectivity, including its cost-effective and non-confining application, there are some limitations that need to be mentioned. First, because fNIRS is limited in monitoring activity from (superficial) cortical structures (Lloyd-Fox et al., 2010), our approach did not allow us to measure activity from deeper cortical and subcortical regions and include those in our network analyses. Second, from a developmental perspective, it should be noted that our analysis is limited to only one age group and comprised of very young infants. It is thus critically important to further assess the development of variability in these brain networks and their associations with behavioral temperament over developmental time to determine its long-term effects and the robustness of these associations (Imai et al., 2014).

In conclusion, the current study provides novel insights into the use of fNIRS in identifying neural endophenotypes -variability in functional brain network connectivity-linked to behavioral temperament traits in early human development. The present findings support the notion that functionally distinct neural networks are implicated in regulatory and emotional behaviors already in newborn infants, adding a critical developmental component to efforts directed at mapping how the individual functional connectome links to affective, cognitive and behavioral traits. The current findings shed light on the brain origins of individual differences in early-emerging behavioral traits and provide the basis for future research examining the genetic and environmental factors contributing to and the long-term developmental consequences of this brain-behavior correlation. More generally, the current study provides early ontogenetic evidence for the idea that studying functional brain network connectivity is an effective way in helping bridge the gap between brain and behavior.

## Acknowledgements

We are grateful to all families who participated in this study as well as Sarah Thomas, Christina Marlow, Kate Haynes, Carolynn McElroy, Julia Larsen, Heath Yancey, Sujal Sigdel, and Shefalika Prasad for assistance with infant data collection at the University of Virginia. This research was supported by *Danone North America, Gut Microbiome, Yogurt and Probiotics Fellowship Grant, Jefferson Scholars Foundation* and *UVA Data Science Fellowship* (to C.M.K) and National Science Foundation #2017229 and *UVA Brain Institute Seed fund* (to T.G).

## Preprint

This manuscript is available as a pre-print on BioRxiv (link: https://www.biorxiv.org/content/10.1101/2020.07.15.204271v1)

## Data Availability

Data are available upon request to the authors.

