## Supplementary Materials for "Variability in infants’ functional brain network connectivity is associated with differences in affect and behavior"

Caroline M. Kelsey^1*^, Katrina Farris^1^ & Tobias Grossmann^1,2^

^1^ Department of Psychology, University of Virginia, Charlottesville, VA, USA

^2^ Max Planck Institute for Human Cognitive and Brain Sciences, Leipzig, Germany

^*^Correspondence concerning this article should be addressed to:

Caroline Kelsey

Department of Psychology

PO BOX 400400

University of Virginia

Charlottesville, VA 22904

Tobias Grossmann

**Descriptive statistics for channels**

*Supplementary Table 1*. Descriptive statistics for functional connections across networks. Note, associations that survive bonferonni corrections are in bold.

|  |  | oxyHb | | | deoxyHb | | |
| --- | --- | --- | --- | --- | --- | --- | --- |
| Channels | N | Mean | SD | Sig. | Mean | SD | Sig. |
| HIN |  |  |  |  |  |  |  |
| 1--26 | 62 | **0.16** | **0.36** | **0.001** | 0.16 | 0.50 | 0.017 |
| 2--28 | 65 | **0.28** | **0.46** | **< .001** | **0.17** | **0.41** | **0.001** |
| 3--34 | 71 | **0.45** | **0.56** | **< .001** | **0.33** | **0.56** | **< .001** |
| 4--29 | 53 | 0.06 | 0.28 | 0.149 | 0.01 | 0.23 | 0.65 |
| 5--31 | 49 | 0.09 | 0.30 | 0.031 | 0.02 | 0.27 | 0.596 |
| 6--38 | 71 | 0.10 | 0.35 | 0.016 | 0.11 | 0.30 | 0.004 |
| 7--32 | 45 | 0.03 | 0.27 | 0.476 | 0.14 | 0.32 | 0.005 |
| 9--35 | 64 | 0.14 | 0.37 | 0.004 | **0.17** | **0.33** | **< .001** |
| 12--30 | 52 | 0.06 | 0.30 | 0.157 | < .001 | 0.24 | 0.921 |
| 13--36 | **71** | **0.29** | **0.48** | **< .001** | **0.23** | **0.40** | **< .001** |
| 14--39 | 68 | 0.12 | 0.35 | 0.006 | 0.12 | 0.37 | 0.01 |
| 15--45 | 62 | 0.07 | 0.32 | 0.106 | 0.14 | 0.39 | 0.006 |
| 16--33 | **71** | **0.26** | **0.36** | **< .001** | **0.24** | **0.38** | **< .001** |
| 17--40 | 72 | 0.10 | 0.33 | 0.01 | 0.11 | 0.33 | 0.007 |
| 18--48 | 68 | 0.11 | 0.40 | 0.02 | 0.06 | 0.37 | 0.161 |
| 19--37 | 70 | 0.01 | 0.26 | 0.677 | 0.02 | 0.34 | 0.593 |
| 20--44 | 62 | 0.07 | 0.33 | 0.1 | 0.03 | 0.32 | 0.487 |
| 21--46 | 59 | 0.07 | 0.32 | 0.116 | -0.05 | 0.30 | 0.226 |
| 22--41 | 67 | 0.07 | 0.29 | 0.052 | 0.05 | 0.36 | 0.25 |
| 23--47 | 64 | 0.05 | 0.30 | 0.202 | 0.02 | 0.32 | 0.638 |
| 24--49 | 57 | 0.08 | 0.35 | 0.104 | 0.08 | 0.37 | 0.119 |
| FPN |  |  |  |  |  |  |  |
| 2--7 | **70** | **0.18** | **0.37** | **< .001** | 0.07 | 0.37 | 0.114 |
| 2--16 | **71** | **0.20** | **0.42** | **< .001** | 0.08 | 0.42 | 0.112 |
| 3--7 | **69** | **0.19** | **0.39** | **< .001** | 0.05 | 0.35 | 0.261 |
| 3--16 | **70** | **0.20** | **0.41** | **< .001** | 0.12 | 0.43 | 0.025 |
| 7--13 | 69 | 0.11 | 0.41 | 0.03 | 0.12 | 0.37 | 0.01 |
| 13--16 | 68 | 0.06 | 0.42 | 0.229 | 0.10 | 0.42 | 0.059 |
| 28--32 | 41 | 0.02 | 0.21 | 0.631 | 0.07 | 0.25 | 0.074 |
| 28--33 | 65 | **0.28** | **0.43** | **< .001** | 0.14 | 0.44 | 0.013 |
| 32--34 | 43 | -0.06 | 0.28 | 0.177 | 0.06 | 0.26 | 0.165 |
| 32--36 | 46 | -0.04 | 0.26 | 0.306 | 0.01 | 0.25 | 0.782 |
| 33--34 | 72 | **0.49** | **0.53** | **< .001** | **0.42** | **0.47** | **< .001** |
| 33--36 | 74 | **0.47** | **0.50** | **< .001** | **0.49** | **0.54** | **< .001** |
| DMN |  |  |  |  |  |  |  |
| 10--18 | 66 | 0.11 | 0.34 | 0.009 | 0.09 | 0.40 | 0.075 |
| 10--22 | 66 | 0.11 | 0.40 | 0.023 | **0.17** | **0.37** | **0.001** |
| 10--23 | 66 | 0.14 | 0.41 | 0.006 | 0.13 | 0.45 | 0.02 |
| 10--24 | 59 | 0.02 | 0.38 | 0.685 | 0.09 | 0.31 | 0.029 |
| 10--41 | 61 | 0.08 | 0.41 | 0.158 | 0.09 | 0.44 | 0.104 |
| 10--47 | 60 | 0.11 | 0.42 | 0.046 | 0.01 | 0.45 | 0.89 |
| 10--48 | 63 | 0.10 | 0.41 | 0.053 | 0.03 | 0.47 | 0.57 |
| 10--49 | 62 | 0.11 | 0.37 | 0.025 | 0.12 | 0.33 | 0.006 |
| 18--42 | 69 | 0.08 | 0.32 | 0.036 | 0.03 | 0.34 | 0.512 |
| 18--43 | 65 | -0.01 | 0.35 | 0.78 | 0.03 | 0.33 | 0.499 |
| 22--42 | 70 | 0.02 | 0.31 | 0.661 | 0.08 | 0.35 | 0.056 |
| 22--43 | 65 | 0.08 | 0.32 | 0.048 | 0.07 | 0.34 | 0.084 |
| 23--42 | 69 | 0.05 | 0.27 | 0.107 | 0.05 | 0.31 | 0.193 |
| 23--43 | 64 | 0.07 | 0.30 | 0.053 | 0.09 | 0.37 | 0.067 |
| 24--42 | 60 | 0.14 | 0.34 | 0.003 | 0.09 | 0.36 | 0.052 |
| 24--43 | 57 | 0.05 | 0.33 | 0.264 | 0.08 | 0.35 | 0.09 |
| 41--42 | 67 | **0.55** | **0.54** | **< .001** | **0.36** | **0.56** | **< .001** |
| 41--43 | 62 | **0.41** | **0.61** | **< .001** | **0.40** | **0.60** | **< .001** |
| 42--47 | 63 | **0.23** | **0.40** | **< .001** | **0.33** | **0.44** | **< .001** |
| 42--48 | 67 | 0.13 | 0.41 | 0.009 | 0.15 | 0.40 | 0.003 |
| 42--49 | 66 | **0.19** | **0.47** | **0.001** | **0.20** | **0.46** | **0.001** |
| 43--47 | 61 | 0.09 | 0.39 | 0.086 | 0.11 | 0.38 | 0.027 |
| 43--48 | 64 | 0.08 | 0.35 | 0.07 | 0.14 | 0.47 | 0.022 |
| 43--49 | 63 | **0.40** | **0.54** | **< .001** | **0.40** | **0.49** | **< .001** |
| Control |  |  |  |  |  |  |  |
| 15--41 | 68 | 0.07 | 0.39 | 0.167 | 0.07 | 0.36 | 0.108 |
| 15--48 | 70 | 0.08 | 0.38 | 0.097 | 0.06 | 0.45 | 0.3 |
| 15--49 | 69 | 0.13 | 0.38 | 0.006 | 0.11 | 0.38 | 0.016 |
| 18--37 | 70 | -0.02 | 0.32 | 0.693 | 0.03 | 0.31 | 0.381 |
| 18--45 | 61 | 0.06 | 0.37 | 0.197 | 0.05 | 0.32 | 0.257 |
| 18--46 | 63 | 0.04 | 0.32 | 0.373 | 0.09 | 0.36 | 0.049 |
| 19--41 | 66 | 0.04 | 0.31 | 0.326 | 0.12 | 0.33 | 0.004 |
| 19--48 | 68 | 0.10 | 0.29 | 0.004 | 0.08 | 0.40 | 0.086 |
| 19--49 | 67 | 0.09 | 0.29 | 0.012 | 0.07 | 0.36 | 0.093 |
| 21--41 | 65 | 0.01 | 0.31 | 0.779 | -0.03 | 0.30 | 0.448 |
| 21--48 | 66 | 0.02 | 0.31 | 0.532 | 0.03 | 0.37 | 0.545 |
| 21--49 | 65 | 0.06 | 0.30 | 0.138 | 0.09 | 0.32 | 0.032 |
| 22--37 | 70 | 0.02 | 0.30 | 0.648 | 0.00 | 0.30 | 0.911 |
| 22--45 | 60 | **0.13** | **0.29** | **0.001** | **0.13** | **0.32** | **0.002** |
| 22--46 | 62 | 0.07 | 0.24 | 0.033 | -0.02 | 0.33 | 0.657 |
| 24--37 | 60 | 0.03 | 0.39 | 0.614 | 0.02 | 0.26 | 0.527 |
| 24--45 | 53 | -0.01 | 0.33 | 0.795 | 0.10 | 0.28 | 0.015 |
| 24--46 | 54 | 0.12 | 0.35 | 0.012 | 0.03 | 0.28 | 0.421 |

**Amount of time included and connectivity levels.**

Correlation analyses were conducted to assess the possible relation between the amount of useable data and connectivity in each of the networks. Here, we found a significant positive correlation between amount of useable data and connectivity (oxyHb) in the FPN, *r* = .23, *p* = .043. There were no other significant associations found (all other *p-values* < .20). Additional models were run with total amount of useable data as a covariate for any models that had significant results for the FPN; however, all results remained the same.

**Covariate Associations**

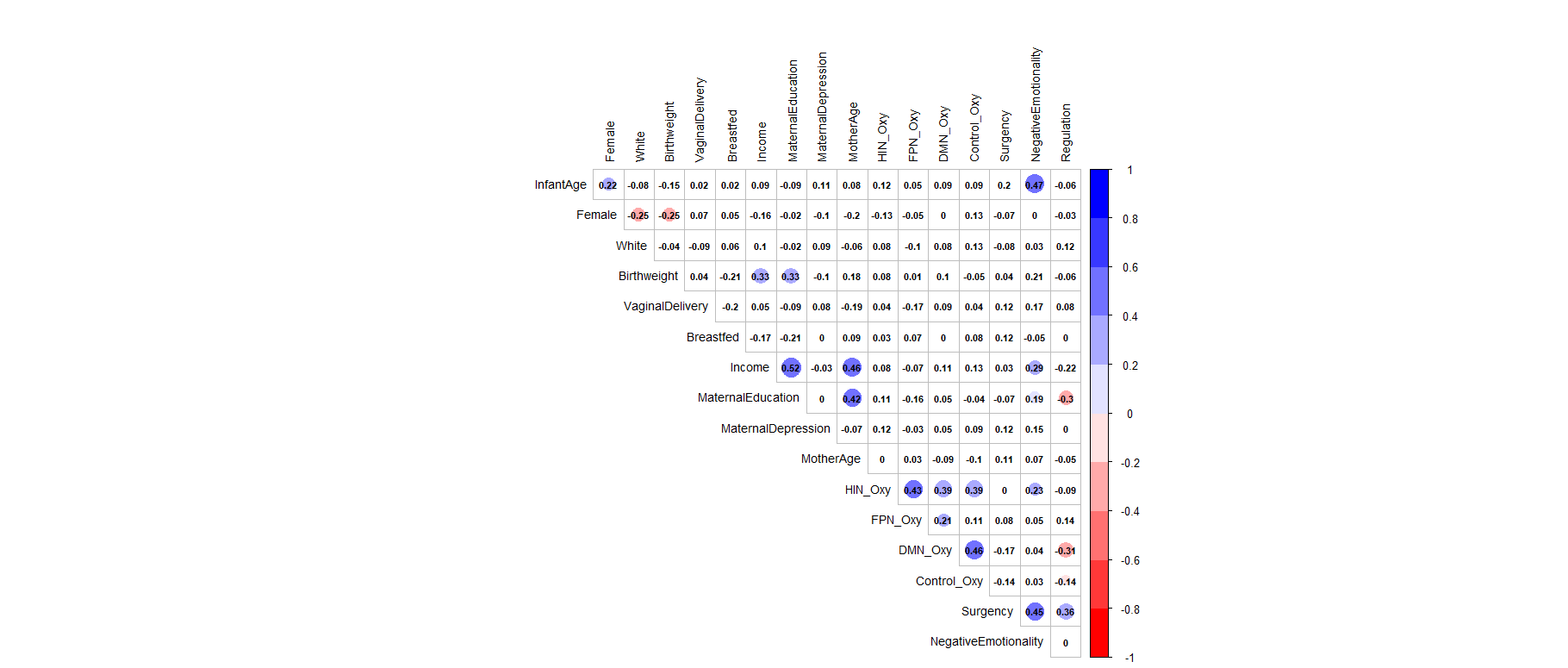

*Supplementary Figure 1.* Spearman’s correlations between socio-demographic factors and study variables. Note, significant associations (*p* < .05) are marked by having a colorful circle in the background.

**Correlations between networks (OxyHb and DeoxyHb)**

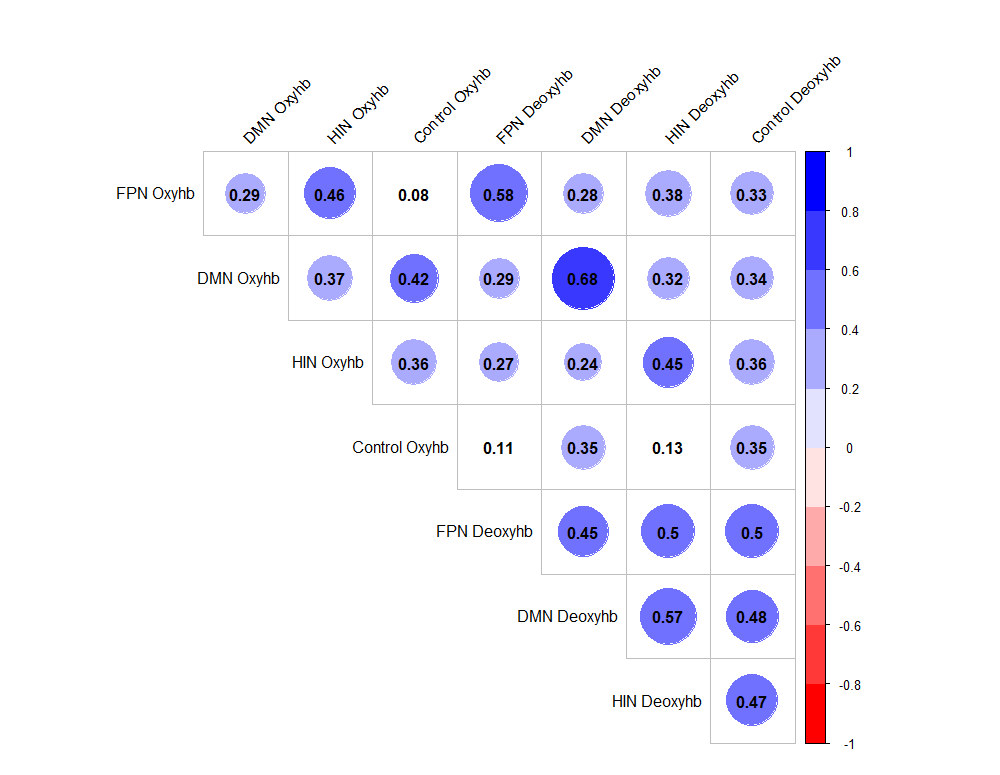

*Supplementary Figure 2.* Correlations between functional networks. Note, significant associations (*p* < .05) are marked by having a colorful circle in the background.

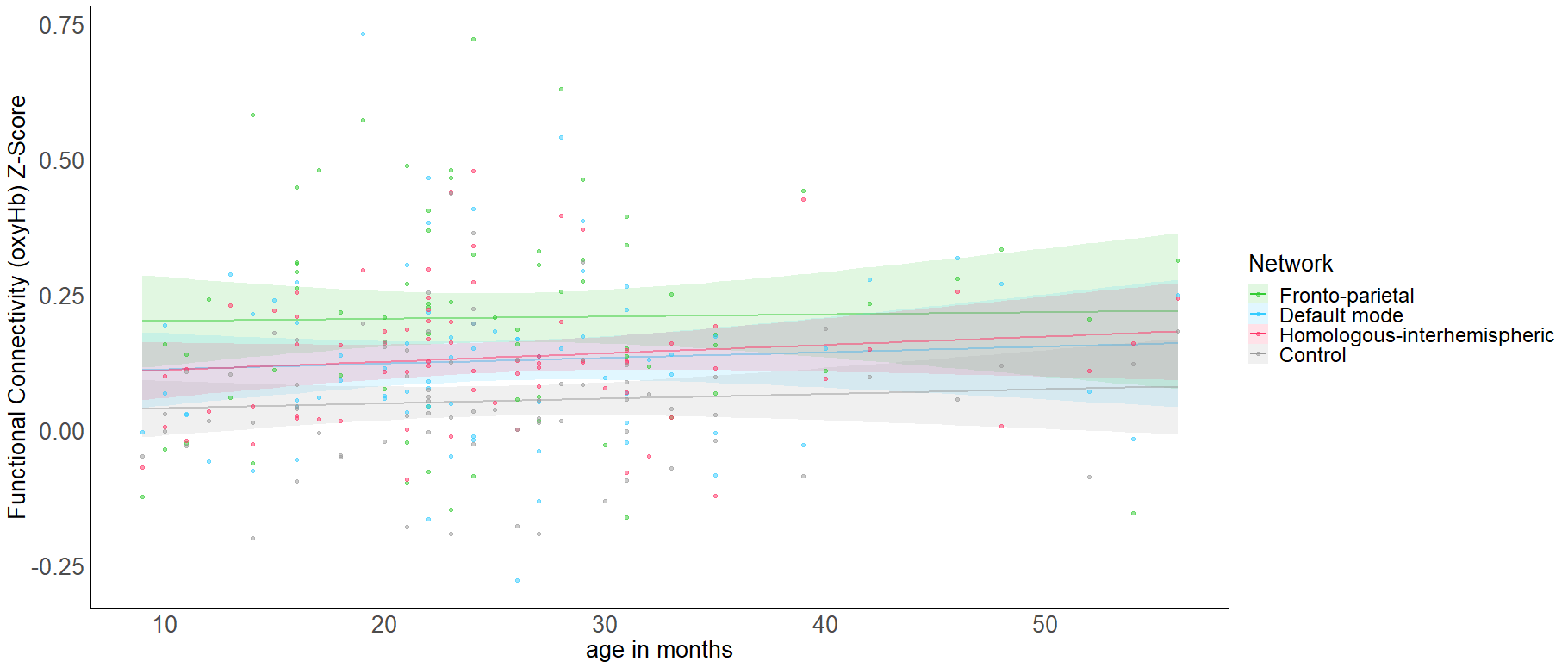

*Supplementary Figure 3.* Depiction of connectivity ranges by infants’ age in months. Note, there are no significant associations between age and functional connectivity levels in any of the networks.

**DeoxyHb Results**

**Functional connectivity (deoxyHb) across networks.**

To analyze differences in overall connectivity levels across networks for deoxyHb an omnibus repeated measures ANOVA with network type (homologous-interhemispheric network [HIN], default mode network [DMN], fronto-parietal network [FPN], control) as a within-subjects factor was conducted. This analysis revealed a significant within-subjects effect across network types, *F*(3, 222) = 11.77, *p* < .001, η^2^ = .137. Post-hoc analyses with Bonferroni adjustments for multiple comparisons were conducted to assess which networks significantly differed from one another. Here, we found that control network (*M* = .05; *SD* = .12) had significantly lower connectivity than the FPN (*M* = .16; *SD* = .20), p < .001, HIN (*M* = .11; *SD* = .12), *p* = .001, and the DMN (*M* = .13; *SD* = .16), *p* < .001. However, there was no significant difference found between the functional networks of interest, all *p’s >* .08 (see *Supplementary Figure 4* for more information).

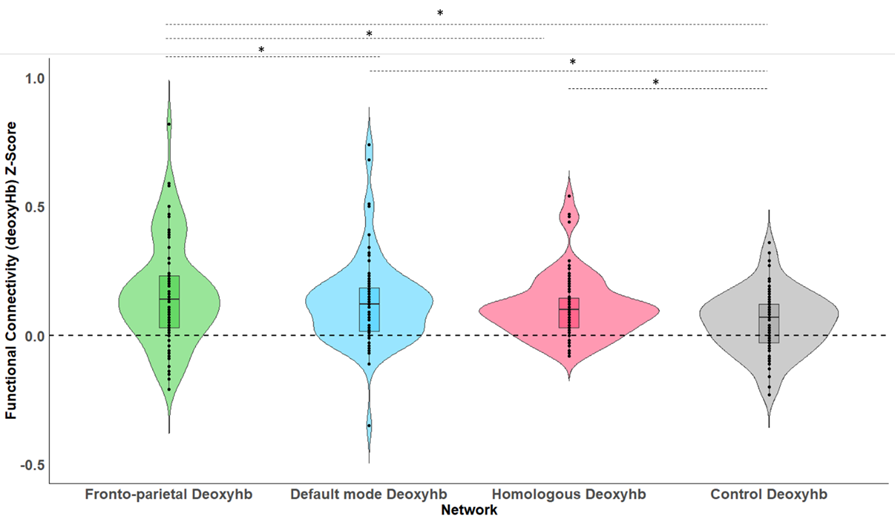

*Supplementary Figure 4.* Shows the average levels of functional connectivity (deoxyHb) and range of variability for each network. The boxplot horizontal lines from bottom to top reflect values for the lower quartile, median, and upper quartile respectively. Note, * *p* < .05.

**Functional connectivity (deoxyHb) and temperament.**

In order to assess how functional connectivity patterns for deoxyHb differentially predicted temperament characteristics, three separate regressions with all four network types (HIN, DMN, FPN, control) predicting each of the three domains of temperament (negative emotionality, regulation/orienting, surgency/positive Emotionality) were conducted.

*Regulation/orienting.* A linear regression was conducted with the four network types (HIN, DMN, FPN, control) predicting regulation/orienting using the entry method. Here, the regression model did not significantly predict regulation/orienting, *p* = .59.

*Negative emotionality.* A linear regression was conducted with the four network types (HIN, DMN, FPN, control) predicting negative emotionality using the entry method. Here, the model did not significantly predict negative emotionality, *p* = .24.

*Surgency/positive emotionality.* A linear regression was conducted with the four network types (HIN, DMN, FPN, control) predicting surgency/positive emotionality using the entry method. Again, the regression model did not significantly predict surgency/positive emotionality, *p* = .67.
